# A Labeled Dataset for AI-based Cryo-EM Map Enhancement

**DOI:** 10.1101/2025.03.16.643562

**Authors:** Nabin Giri, Xiao Chen, Liguo Wang, Jianlin Cheng

**Author notes:** Corresponding author: Jianlin Cheng. these authors contributed equally to this work (Nabin Giri, Xiao Chen).

## Abstract

Cryo-electron microscopy (cryo-EM) has transformed structural biology by enabling near-atomic resolution imaging of macromolecular complexes. However, cryo-EM density maps suffer from intrinsic noise arising from structural sources, shot noise, and digital recording, which complicates accurate atomic structure building. While various methods for denoising cryo-EM density maps exist, there is a lack of standardized datasets for benchmarking artificial intelligence (AI) approaches. Here, we present an open-source dataset for cryo-EM density map denoising comprising 650 high-resolution (1-4 Å) experimental maps paired with three types of generated label maps: regression maps capturing idealized density distributions, binary classification maps distinguishing structural elements from background, and atom-type classification maps. Each map is standardized to 1 Å voxel size and validated through Fourier Shell Correlation analysis, demonstrating substantial resolution improvements in label maps compared to experimental maps. This resource bridges the gap between structural biology and artificial intelligence communities, enabling researchers to develop and benchmark innovative methods for enhancing cryo-EM density maps.

## Background & Summary

Recent advances in cryo-electron microscopy (cryo-EM) have revolutionized structural biology by enabling the visualization of macromolecular complexes at near-atomic resolution^1,2^. Cryo-EM density maps are the three-dimensional volume reconstructions derived from two-dimensional images captured by electron microscopy. They have become invaluable tools for understanding complex biological structures. However, the experimental process inherently introduces artifacts that affect image quality and subsequent interpretability.

The generation of cryo-EM density maps involves the rapid exposure of flash-frozen aqueous samples to electron beams in vacuum. This process inherently produces raw images with low signal-to-noise ratios (SNR), significantly complicating the accurate interpretation of structural details. The resulting three-dimensional reconstructions consequently suffer from noise-related challenges that obstruct precise and accurate determination of atomic arrangements within the density maps. Given that the ultimate goal of cryo-EM single particle analysis is to obtain accurate atomic structures of protein complexes, these noise-related limitations present substantial obstacles to the structure building process.

Noise in cryo-EM data manifests at three distinct stages: First, *structural noise* arises from the surrounding ice matrix and often a superimposed thin carbon film. This background structure varies from one molecule to the next and is therefore inherently irreproducible. Conceptually, *structural noise* also encompasses any conformational variations within the molecule itself that are not consistently reproduced across samples. Second, *shot noise* results from the quantum nature of electron radiation, introducing statistical variations in electron detection. Third, *digital noise* emerges during the recording and digitization process whether from photographic granularity and microdensitometer digitization noise in traditional methods, or from readout noise in modern direct electron detectors. While *shot* and *digital* noise appear as random, dust-like distributions without specific patterns (background noise), *structural noise* exhibits defined shapes with stronger density signals that can particularly confound interpretation^3^.

The combined effect of these noise sources pass on through every stage of the structural determination pipeline, from initial data collection to final 3D reconstruction of cryo-EM density maps. This noise accumulation significantly hampers the global and local resolution and interpretability of the resulting density maps, making accurate atomic model building challenging. In particular, the noise can obscure important features such as side-chain densities and ligand binding sites^4–9^, potentially leading to misinterpretation of structural and functional characteristics^10,11^.

Denoising cryo-EM density maps aims to mitigate these experimental artifacts, significantly enhancing map interpretability by clarifying the three-dimensional arrangement of atoms. This improved clarity is essential for understanding the functional properties of imaged biomolecules. Moreover, noise reduction substantially facilitates downstream atomic structure modeling tasks, benefiting both manual interpretation and automated approaches such as Phenix^12^, MAINMAST^13^, Cryo2Struct^14^, DeepTracer^15^, DeepMainmast^16^, and ModelAngelo^17^. Recent advances in computational methods have shown promise in addressing these challenges, with deep learning approaches demonstrating particular potential for distinguishing signal from noise in complex density distributions^18^.

While various methods have been developed for denoising cryo-EM density maps, the field lacks standardized, and open-source datasets for advancement of artificial intelligence-based denoising approaches. This deficiency is particularly problematic given the rapid proliferation of AI-based tools in structural biology, as it prevents rigorous comparison between methods and hinders benchmarking of new approaches. Furthermore, the absence of standardized test cases makes it difficult to assess how denoising algorithms perform across diverse structural classes and resolution ranges.

As cryo-EM technology continues its rapid expansion in structural biology, with applications extending to increasingly complex systems such as membrane proteins, large macromolecular assemblies, and heterogeneous samples, there is a need for benchmark datasets that bridge disciplinary boundaries and enable AI practitioners to develop innovative methods for enhancing map quality and interpretability. The dataset presented in this manuscript addresses this need by providing a comprehensive and standardized resource for the development and evaluation of cryo-EM denoising algorithms. By preparing this dataset, we aim to drive methodological innovation at the intersection of artificial intelligence and structural biology. Our goal is to accelerate progress in cryo-EM density map interpretation and, ultimately, to advance biomedical research through improved macromolecular structure determination from cryo-EM.

## Methods

### Related works

Recently, several AI-based methods have been developed to enhance the quality and interpretability of cryo-EM density maps, including DeepEMhancer^19^, EMReady^20^, DeepTracer-Denoising^21^, and CryoTEN^22^. A key aspect of training these models is the generation of high-quality label maps. Existing approaches include creating simulated cryo-EM density maps using tools like *pdb2mrc* from the EMAN2 package^23^, *pdb2vol* from the Situs package^24^, or methods based on Rosetta^25^. These simulated maps, derived from known atomic structures, serve as idealized, noise-free ground truth representations of protein density. Alternatively, some models are trained on label maps generated with LocScale by leveraging atomic models to refine experimental cryo-EM maps^25^. In this work, we introduce an innovative approach for generating high-quality label maps which, when paired with experimental cryo-EM maps, are used to train AI-based models for denoising and enhancing cryo-EM density maps. Once trained, these models can refine noisy experimental maps, improving their clarity and structural interpretability by reducing artifacts and noise.

### Data Acquisition and Preprocessing

#### Dataset Curation

We curated a dataset of high-resolution cryo-EM density maps for single-particle proteins from the Electron Microscopy Data Bank^26^ (EMDB), selecting those with resolutions between 1 and 4 Å. The corresponding atomic biological assembly structures were retrieved from the Protein Data Bank^27^ (PDB) and used to generate label maps. We refer to the deposited maps obtained from EMDB as experimental cryo-EM density maps, which contain noise and artifacts that must be identified and removed for effective denoising. To ensure data quality, we applied the following filtering criteria: (1) Removal of maps without a corresponding atomic biological assembly structure in the PDB. (2) Removal of maps missing a resolution value determined by the Fourier Shell Correlation (FSC) 0.143 cut-off score. (3) Removal of redundant cryo-EM density maps associated with the same atomic biological assembly structure.

The FSC was computed using the *phenix*.*mtriage* function from Phenix software suite^12^ by providing the experimental cryo-EM density map and its associated PDB structure as input. After filtering, we obtained a final dataset of 650 cryo-EM density maps, which we used to generate labels for training, validation, and testing of deep learning models for cryo-EM density map denoising.

#### Experimental Map Standardization

Experimental cryo-EM density maps in EMDB exhibit significant variations in density values (e.g., [-2.32, 3.91] and [-0.553, 0.762]) and voxel sizes (e.g., ranging from 0.7 Å to 1.6 Å) due to differences in microscope models, electron doses, detectors, and imaging conditions. Since our approach aims to refine cryo-EM density maps at the voxel level through voxel-wise classification or regression models, we standardized all experimental maps to ensure consistency. We standardized the voxel size of all experimental cryo-EM density maps to 1 Å using the resampling function in UCSF ChimeraX^28^. This preprocessing step enables the model to learn meaningful patterns across different cryo-EM density maps, ultimately improving its robustness and accuracy in denoising and structure enhancement.

### Label Map Generation Workflow

We generated three types of label maps to train AI-based models for cryo-EM density map enhancement: a regression map capturing density values representing an ideal, noise-free cryo-EM map, and two classification label maps that provide complementary information to improve model learning during training.

#### Simulated Map Generation

In the first stage of label map generation, we created clean, noise-free cryo-EM density maps from atomic biological PDB structures using the *pdb2vol* utility from the Situs package^24^. This real-space convolution software generates simulated volumetric maps from input atomic structures. The simulated maps were generated with a voxel size of 1 Å, ensuring consistency with the standardized experimental cryo-EM density maps.

#### Label Map Generation

While simulated maps provide an idealized, artifact-free representation of cryo-EM density, they often lack precise voxel-wise alignment with their corresponding experimental maps. This alignment is important for computing accurate voxel-wise errors during model optimization. To address this, we processed the simulated maps to ensure they match the size, voxel spacing, and voxel-wise alignment of the experimental maps. We created three empty mask maps (meaning, the voxel values are filled with zero), each matching the dimensions of the experimental cryo-EM density map, to store label information. Using atomic coordinates from the PDB structure, we converted Euclidean coordinates into voxel grid indices using the Formula 1, ensuring proper mapping between the experimental and simulated maps. To address the challenges in label generation, we created three distinct types of label maps:

1. **Regression Label Map:** This map contains continuous density values derived from the simulated cryo-EM density map. At each voxel position corresponding to an atom in the PDB structure, we assign the density value from the simulated map, providing a noise-free target for regression-based learning.
2. **Classification Label Map:** This binary classification map distinguishes structural elements from background. We assign a value of 1 to voxels corresponding to atomic positions while background voxels remain at 0.
3. **Atom Type Classification Label Map:** This multi-class map differentiates between atom types, enabling the model to learn chemical specificity. We assign distinct values to voxels based on the corresponding atom type: C*α* (1), C*β* (2), carbonyl carbon (3), oxygen (4), and nitrogen (5).

A critical consideration in our label generation approach is the treatment of neighboring voxels. Due to the inherent discrepancy between continuous atomic coordinates and discrete voxel grid positions, simply labeling the exact voxel corresponding to each atom would result in sparse and potentially imprecise labels. Furthermore, electron density in cryo-EM maps extends beyond the exact atomic positions, forming a cloud-like distribution that represents the probability of electron occurrence.

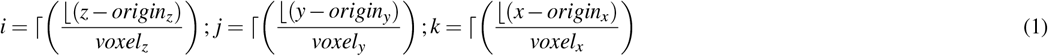

To address these issues and improve the structural context, we extended the labeled regions by identifying neighboring voxels within a 6 Å radius of each atomic coordinate. This radius was selected based on analysis of electron density distribution in high-resolution cryo-EM maps. This is necessary to enhance interpretability by providing meaningful structural context beyond atomic locations and to minimize precision loss when converting atomic coordinates (continuous space) into voxel grid indices (discrete space) as shown in Formula 1. The approach for handling these neighboring voxels varies by label map type:

- For the **Regression Label Map**, neighboring voxels are assigned corresponding values from the simulated cryo-EM density map, creating a continuous gradient of density values that mimics the electron cloud surrounding each atom.
- For the **Classification Label Map**, neighboring voxels are assigned a distinct value of 2, differentiating them from both the central atomic positions (value 1) and the background (value 0). This three-class approach allows the model to learn regions between atoms and background.
- For the **Atom Type Classification Label Map**, we maintain the original atomic type labeling only for voxels corresponding directly to atom positions, as the chemical identity of neighboring voxels cannot be unambiguously determined.

By providing complementary information across different label types, we enable deep learning models to learn diverse aspects of molecular structure, ultimately leading to more robust and accurate enhancement of experimental cryo-EM density maps. These label maps can be used individually or in combination during model training, allowing for flexible architecture design and task-specific optimization strategies.

### Data Records

Our dataset consists of the following components, curated to support the development and validation of deep learning models for cryo-EM density map enhancement:

#### Experimental Cryo-EM Density Maps

We compiled 650 high-resolution (1-4 Å) experimental maps from EMDB, standardized to 1 Å voxel size. These maps represent diverse structural classes including soluble proteins, membrane proteins, protein-nucleic acid complexes, and macromolecular assemblies. Each map is stored in MRC format with associated metadata including resolution and PDB accession codes.

#### Atomic Biological Assembly Structures

For each experimental map, we obtained the corresponding biological assembly atomic structure from PDB. These complete assemblies ensure full occupancy of the experimental density maps and serve as the foundation for both label generation and quality validation.

#### Simulated Cryo-EM Density Maps

Using the *pdb2vol* utility from Situs, we generated idealized, noise-free density maps from the atomic structures. These simulated maps serve as intermediates in our label generation process and are created with parameters matched to their experimental counterparts.

#### Label Maps

Based on the atomic structures and simulated maps, we generated three types of label maps:

- Regression Label Maps: Contain continuous density values derived from simulated maps, spatially aligned to the experimental maps. These maps represent idealized electron density distributions and serve as primary training targets.
- Classification Label Maps: Differentiate protein structure from background using a three-value system: atomic positions (1), neighboring regions within 6 Å (2), and background (0).
- Atom Type Classification Label Maps: Label chemical information by assigning values based on atom type: C*α* (1), C*β* (2), carbonyl carbon (3), oxygen (4), and nitrogen (5).

All data records maintain consistent dimensions and spatial alignment with their corresponding experimental maps, ensuring proper voxel-to-voxel correspondence for training deep learning models in supervised learning manner. The dataset is organized by PDB identifiers, with supplementary metadata files providing resolution information and quality metrics.

### Technical Validation

To objectively validate the quality of our regression label maps compared to the experimental maps, we used Fourier Shell Correlation (FSC), the gold standard for resolution assessment in cryo-EM. FSC quantifies the normalized cross-correlation between two 3D volumes as a function of spatial frequency, providing an objective measure of similarity between maps and their corresponding atomic models. For each map in our dataset (n = 650), we computed FSC using the *phenix*.*mtriage* tool from the Phenix software suite^12^. We analyzed both the standard FSC 0.143 criterion (widely accepted as the resolution threshold for cryo-EM maps) and the more stringent FSC 0.5 criterion (indicating higher confidence in map-model agreement), using unmasked analyses to evaluate global map quality.

Figure 1 presents a comprehensive scatter plot comparison of resolution metrics between the deposited experimental maps and our regression label maps. As shown in Figure 1 (a), the FSC 0.143 unmasked resolution values for the regression label maps (orange, mean = 1.95 Å) are better than those for the deposited experimental maps (green, mean = 2.69 Å). This represents a substantial 27.5% improvement in resolution. The clustering of the orange points around the mean indicates consistent quality enhancement across the dataset, with notably fewer outliers compared to the experimental maps. Similarly, Figure 1 (b) illustrates the comparison using the more stringent FSC 0.5 criterion. The mean unmasked FSC 0.5 resolution improved from 4.01 Å for experimental maps to 3.33 Å for regression label maps, demonstrating a 16.9% enhancement. The distribution pattern shows that regression label maps (orange) maintain more consistent quality at this higher confidence threshold, with experimental maps (green) showing variability.

**Figure 1.**
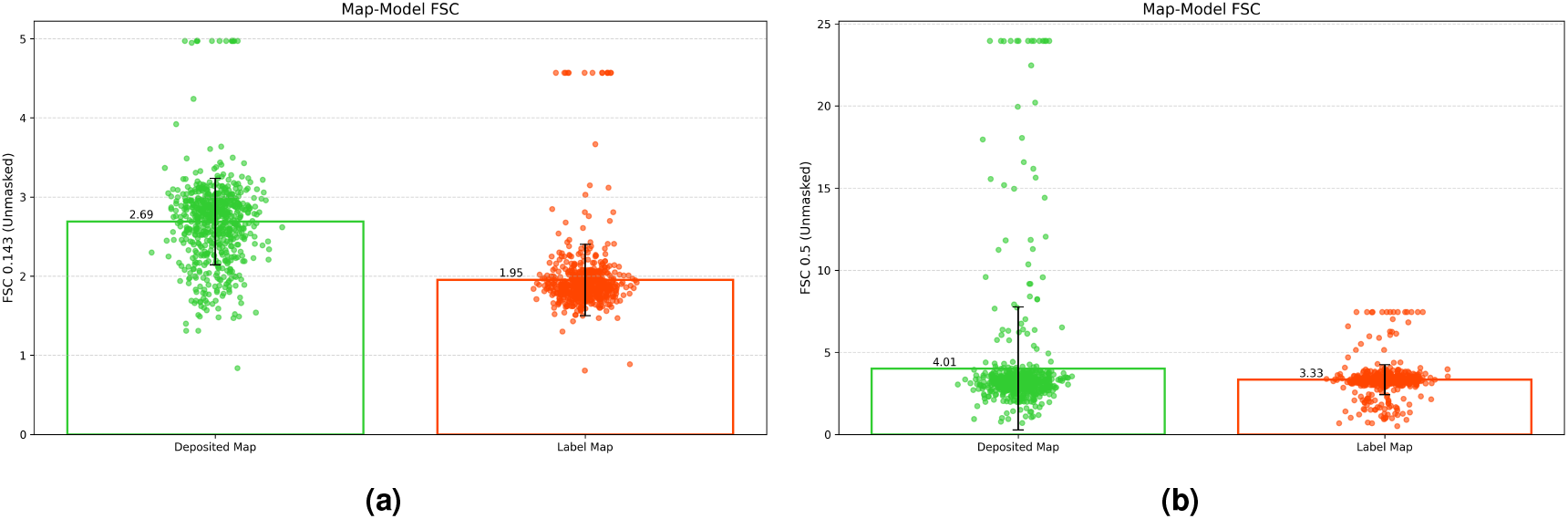
(a) The mean Fourier Shell Correlation (FSC) 0.143 unmasked for deposited map and regression label map is 2.69 Å and 1.95 Å, respectively. (b) The mean FSC 0.5 unmasked for deposited map and regression label map is 4.01 Å and 3.33 Å, respectively.

The box plots in both panels (Figure 1(a) and (b)) highlight the statistical significance of these improvements, with minimal overlap between the distributions. Notably, the upper quartile bound for the regression label maps falls below the mean of the experimental maps in both FSC metrics, emphasizing the consistent better resolution of the label maps.

Figure 2, 3, 4, 5, 6, 7 shows the experimental cryo-EM density map and the regression labeled cryo-EM density map for visual interpretation. These results provide evidence that our regression label maps offer substantially improved structural representation compared to the original experimental maps. The enhanced resolution and consistency make them ideal targets for training deep learning models aimed at denoising and refining experimental cryo-EM density maps. By learning to transform noisy experimental data toward these higher-quality representations, our models can effectively enhance structural interpretability and in turn more accurate de novo atomic modeling.

**Figure 2.**
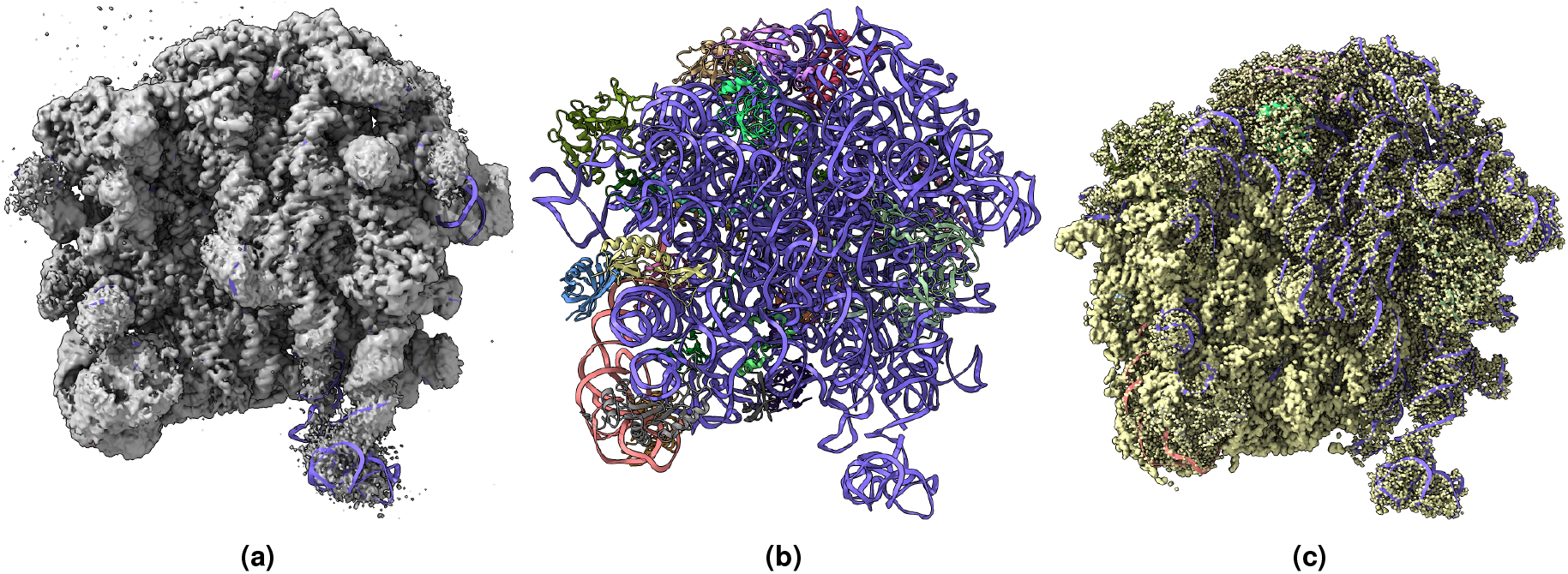
(a) Overlay of the deposited experimental cryo-EM density map (EMD-11900) in grey with dimensions of 308*×* 308 *×*308 and a voxel size of 1*×* 1*×* 1 Å, visualized at the recommended contour level of 0.0037 (1.1 *σ*), along with its corresponding biological atomic structure (PDB Code: 7ASM). The Fourier Shell Correlation (FSC) at 0.5 (unmasked) is 2.43 Å. (b) The atomic structure of the protein (PDB Code: 7ASM). (c) The regression label map in yellow with dimensions of 308 *×*308 *×*308 and a voxel size of 1 *×*1*×* 1 Å, overlaid with the known biological atomic structure (PDB Code: 7ASM), with an FSC at 0.5 (unmasked) of 1.27 Å.

**Figure 3.**
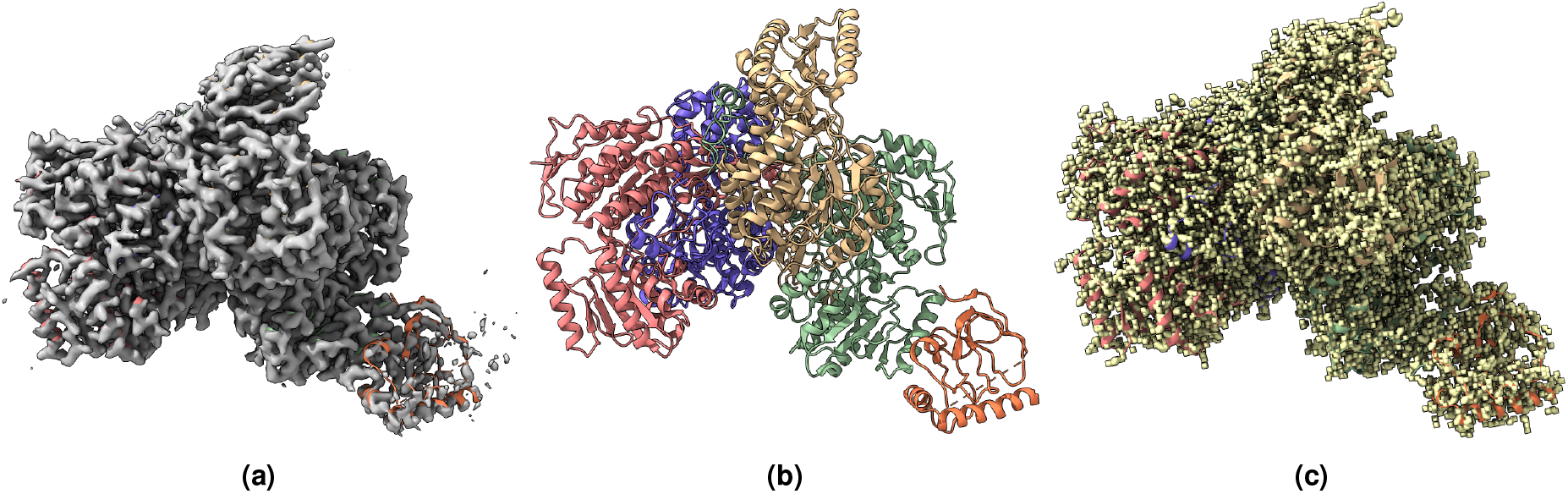
(a) Overlay of the deposited experimental cryo-EM density map (EMD-33113) in grey with dimensions of 283 *×*283*×* 283 and a voxel size of 1 *×*1 *×*1 Å, visualized at the recommended contour level of 0.0178 (4.9 *σ*), along with its corresponding biological atomic structure (PDB Code: 7XC6). The Fourier Shell Correlation (FSC) at 0.5 (unmasked) is 4.16 Å. (b) The atomic structure of the protein (PDB Code: 7XC6). (c) The regression label map in yellow with dimensions of 283 *×*283 *×*283 and a voxel size of 1*×* 1*×* 1 Å, overlaid with the known biological atomic structure (PDB Code: 7XC6), with an FSC at 0.5 (unmasked) of 3.24 Å.

**Figure 4.**
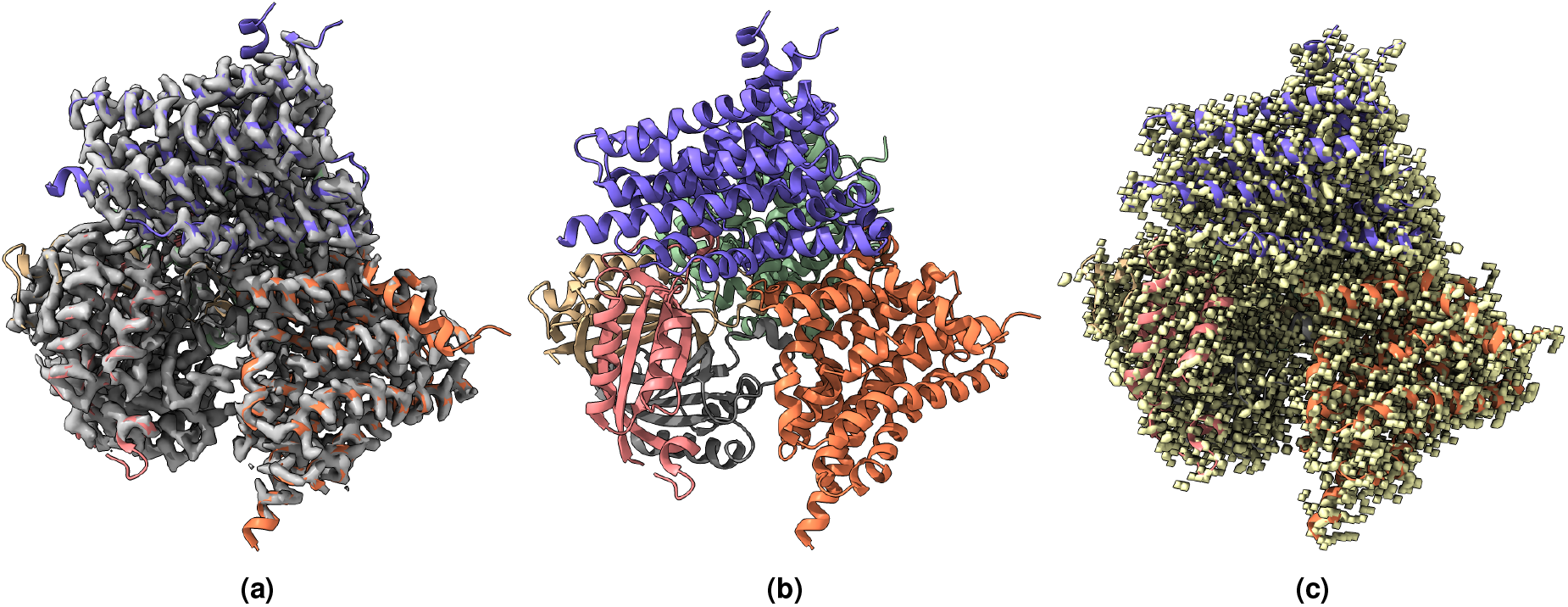
(a) Overlay of the deposited experimental cryo-EM density map (EMD-31135) in grey with dimensions of 258 *×*258 *×*258 and a voxel size of 1*×* 1 *×*1 Å, visualized at the recommended contour level of 0.05 (11.5 *σ*), along with its corresponding biological atomic structure (PDB Code: 7EGK). The Fourier Shell Correlation (FSC) at 0.5 (unmasked) is 7.93 Å. (b) The atomic structure of the protein (PDB Code: 7EGK). (c) The regression label map in yellow with dimensions of 258 *×*258*×* 258 and a voxel size of 1*×* 1 *×*1 Å, overlaid with the known biological atomic structure (PDB Code: 7EGK) with an FSC at 0.5 (unmasked) of 3.23 Å.

**Figure 5.**
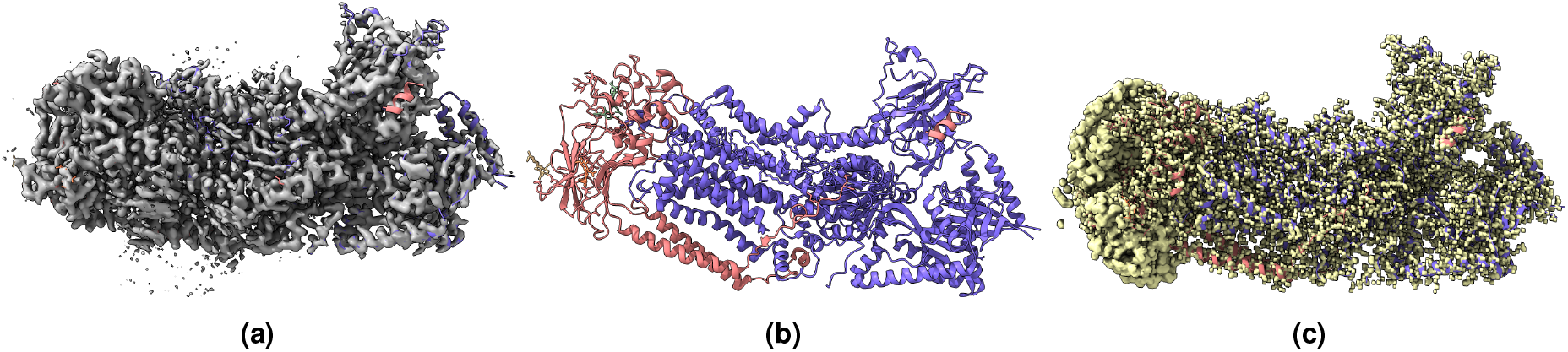
(a) Overlay of the deposited experimental cryo-EM density map (EMD-23075) in grey with dimensions of 231*×* 231*×* 231 and a voxel size of 1 *×*1 *×*1 Å, visualized at the recommended contour level of 0.018 (4.3 *σ*), along with its corresponding atomic structure (PDB Code: 7KYC). The Fourier Shell Correlation (FSC) at 0.5 (unmasked) is 2.86 Å. (b) The atomic structure of the protein (PDB Code: 7KYC). (c) The regression label map in yellow with dimensions of 231 *×*231 *×*231 and a voxel size of 1 *×*1 *×*1 Å, overlaid with the known biological atomic structure (PDB Code: 7KYC) with an FSC at 0.5 (unmasked) of 1.56 Å.

**Figure 6.**
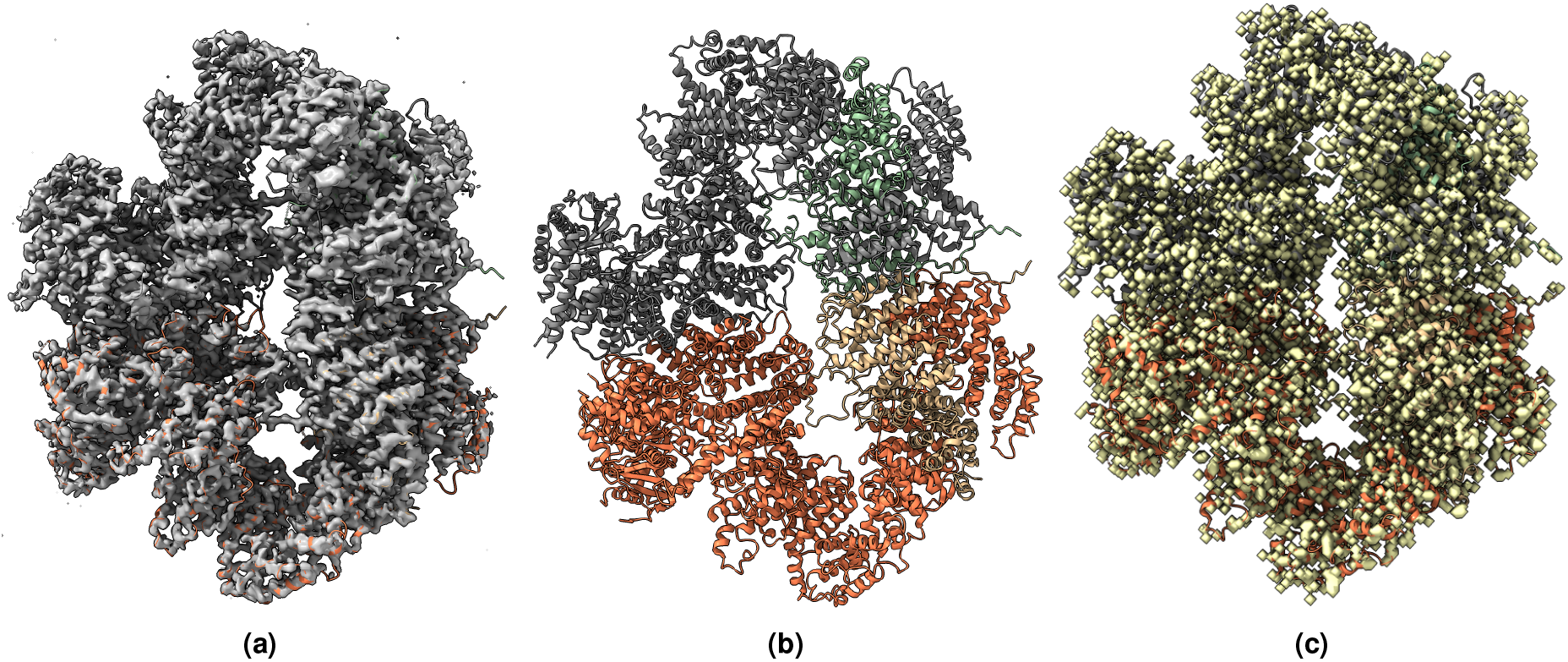
(a) Overlay of the deposited experimental cryo-EM density map (EMD-11055) in grey with dimensions of 347*×* 347*×* 347 and a voxel size of 1 *×*1*×* 1 Å, visualized at the recommended contour level of 1.6 (4.9 *σ*), along with its corresponding biological atomic structure (PDB Code: 6Z2W). The Fourier Shell Correlation (FSC) at 0.5 (unmasked) is 6.32 Å. (b) The atomic structure of the protein (PDB Code: 6Z2W). (c) The regression label map in yellow with dimensions of 347 *×*347 *×*347 and a voxel size of 1*×* 1 *×*1 Å, overlaid with the known biological atomic structure (PDB Code: 6Z2W) with an FSC at 0.5 (unmasked) of 3.32 Å.

**Figure 7.**
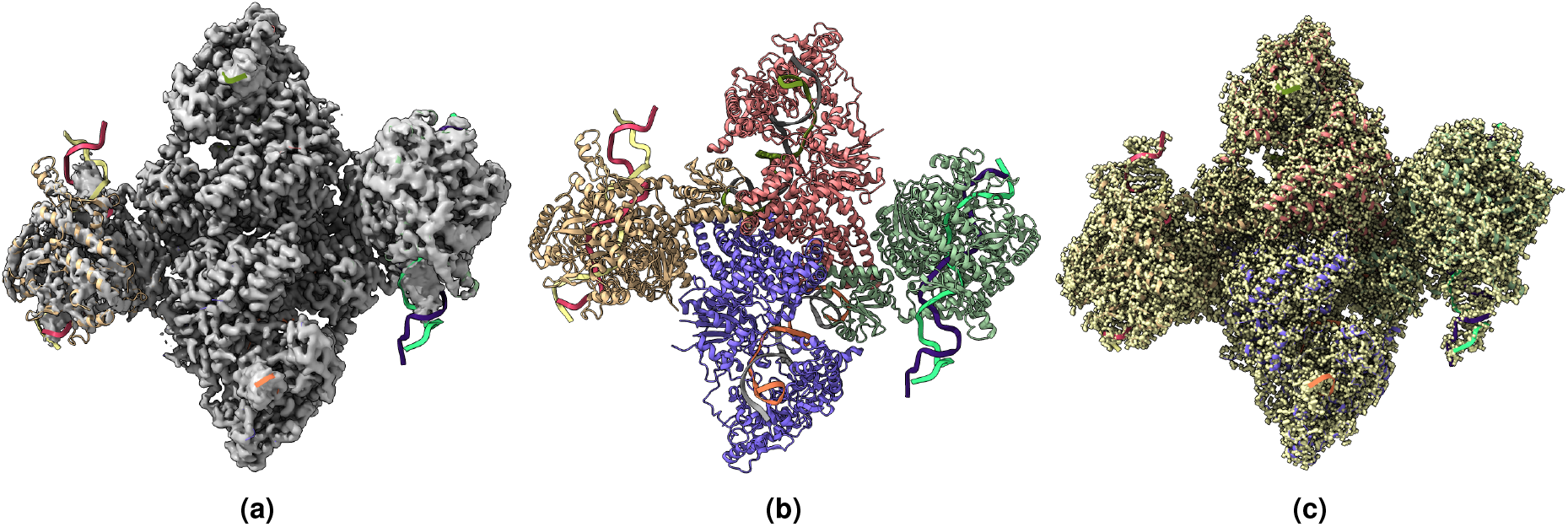
(a) Overlay of the deposited experimental cryo-EM density map (EMD-23461) in grey with dimensions of 526*×* 526 *×*526 and a voxel size of 1*×* 1*×* 1 Å, visualized at the recommended contour level of 0.18 (8.2 *σ*), along with its corresponding biological atomic structure (PDB Code: 7LO5). The Fourier Shell Correlation (FSC) at 0.5 (unmasked) is 6.52 Å. (b) The atomic structure of the protein (PDB Code: 7LO5). (c) The regression label map in yellow with dimensions of 526*×* 526 *×*526 and a voxel size of 1 *×*1*×* 1 Å, overlaid with the known biological atomic structure (PDB Code: 7LO5) with an FSC at 0.5 (unmasked) of 3.5 Å.

### Usage Notes

The dataset provided in this work is specifically designed for training and evaluating deep learning models aimed at improving the clarity and interpretability of cryo-EM density maps. By enhancing cryo-EM map quality, these models, in turn, facilitate more accurate 3D atomic structure modeling from experimental cryo-EM density maps^18^.

The dataset supports the development of various deep learning models for denoising cryo-EM density maps. It can be used for a supervised denoising approach by leveraging pairs of experimental cryo-EM density maps and regression labels to remove noise while preserving structural features. The classification labels allow for semantic segmentation, enabling models to distinguish protein structural components from the background. Additionally, the atom-type classification labels can be used to develop models capable of identifying specific atoms in the cryo-EM density maps.

For optimal results when training deep learning models with this dataset, we recommend that users split the dataset based on resolution, as described in Cryo2StructData^29^, rather than using a random split. To improve memory efficiency, an alternative approach is to generate 3D sub-cubes (e.g., 64 *×*64 *×*64) from the cryo-EM maps instead of processing entire volumes at once. We also encourage users to consider a multi-task learning approach, training on multiple label types to enhance feature learning and generalization.

By following these guidelines, researchers can effectively leverage this dataset to develop robust deep learning models for cryo-EM map enhancement, ultimately advancing structural biology research through improved map interpretation and atomic modeling.

## Code availability

The source code and the instructions to reproduce the dataset is provided in the GitHub repository accessible at https://github.com/BioinfoMachineLearning/denoisecryodata.git. To keep the generated data files permanent, we published all data to the Harvard Dataverse (https://doi.org/10.7910/DVN/CI0J2B), an online data management and sharing platform with a permanent Digital Object Identifier number.

## Examples

## Acknowledgments

This work was supported in part by an NIH grant (R01GM146340). We would like to thank Dr. Liguo Wang for insightful comments.

## Author contributions statement

JC conceived the experiment. JC, NG and XC designed the experiment. NG and XC conducted the experiment. NG, XC, JC and LW analyzed the results. NG and XC drafted the manuscript. NG, XC and JC edited the manuscript.

